# Spatial distribution of cytoskeleton-mediated feedback controls cell polarization: a computational study

**DOI:** 10.1101/2025.04.12.648264

**Authors:** Parijat Banerjee, Jonathan A. Kuhn, Dhiman Sankar Pal, Yu Deng, Tatsat Banerjee, Peter N. Devreotes, Pablo A. Iglesias

## Abstract

In the social amoeba *Dictyostelium*, cell motility is regulated through a signal transduction excitable network that interfaces with the cytoskeleton to control actin polymerization patterns. In turn, the cytoskeleton influences the signaling machinery via several feedback loops, but the nature and function of this feedback remain poorly understood. In this study, we use computational models to discern the essential role of complementary positive and negative feedback loops in polarizing cells. We contrast two potential mechanisms for the negative feedback: local inhibition and global inhibition. Our results indicate that both mechanisms can stabilize the leading edge and inhibit actin polymerization in other sites, preventing multipolarity. While some experimental perturbations align more closely with the local inhibition model, statistical analyses reveal its limited polarization potential within a wide excitability range. Conversely, global inhibition more effectively suppresses secondary and tertiary leading-edge formation, making it a more robust polarization mechanism. This raises an intriguing question: if local inhibition better replicates experimental observations but is less effective for polarization than local excitation and global inhibition, could there be a supplementary mechanism enhancing its polarization potential? To address this, we propose a novel mechanism involving the dynamic partitioning of back molecules which enhances communication between the front and back of the cell and can be leveraged by local feedback interactions between the cytoskeleton and signal transduction to improve polarization efficiency.

## INTRODUCTION

Directed cell motility, the ability of motile cells to sense external cues and use these to navigate, is a fundamental process in various biological contexts, including embryonic development, wound healing, and immune responses. To migrate, cells must develop a spatial asymmetry consisting of well-defined front and rear components. Proteins that localize to either region can be described as front or back markers. The former includes signaling proteins such as Ras and PI3K that regulate actin polymerization, as well as cortical proteins that give rise to protrusions, including actin and actin-binding proteins such as cofilin, coronin, or Arp2/3. Similarly, the rear is marked by various signaling markers (e.g. PI(4,5)P_2_, PTEN, PIP5K), as well as cortical proteins such as myosin II and cortexillin that provide contractility.

Cell polarity can arise in response to spatially heterogeneous chemical or mechanical cues from the environment. These external signals induce internal asymmetries that are temporal — once the cue is removed, the cell reverts to its pre-stimulus, unpolarized state. For example, latrunculin-treated *Dictyostelium* cells, which have inhibited actin polymerization, form crescents of intracellular markers aligned with a chemoattractant gradient despite their impaired motility [1, 2]. However, when the chemoattractant is removed, these cells lose their asymmetry and reorient homogeneously. Alternatively, symmetry breaking can result in a polarized state that persists even after the stimulus is withdrawn; for instance, *Dictyostelium* cells moving toward a gradient maintain their directionality after the cue is removed [3]. To distinguish between these processes, we refer to stable spatial asymmetries as *polarization*, while transient responses to spatially heterogeneous stimuli are described as gradient sensing [3]. Here, we focus primarily on polarization. Note that cells can also spontaneously polarize from stochastic spatial heterogeneities that become amplified and stabilized.

In parallel to the polarization process, cell motility involves dynamic spatial patterning driven by the interaction of two excitable systems: the signal transduction excitable network (STEN) and the cytoskeleton excitable network (CEN). Although these systems are coupled, both exhibit excitability independently when the other is eliminated [4]. It is well-established that the signaling patterns generated by STEN coordinate the actions of CEN to drive migration. There is also substantial evidence that feedback from CEN also influences STEN [5]. While actin polymerization provides positive feedback to signaling at the front of the cell [2, 6–17], at the rear, myosin is involved in distinct feedback loops [18–20]. Ras/PI3K activation promotes myosin disassembly [21–23], and recent findings in *Dictyostelium* show that myosin assembly inhibits Ras/PI3K activation [24]. These findings suggest the presence of a double negative feedback loop — functionally equivalent to positive feedback — that operates at the rear of migrating cells.

In this paper, we explore the interaction between these two patterning phenomena to understand how feedbacks arising from coupled excitable systems can stabilize to form polarized cells (Fig. 1). To achieve this, we rely on computational modeling. The study of spatial patterns has long benefited from mathematical models, beginning with Alan Turing’s pioneering concept of morphogenesis—the development of spatial patterns from an initially homogeneous state [25]. Turing’s work showed how differential diffusion properties between activating and inhibiting processes could destabilize homogeneous states, leading to the emergence of patterns. Building on this, Meinhardt and Gierer formalized a model based on local self-enhancing activation and lateral inhibition (LALI) [26]. The minimal LALI model proposes two interacting species: a local activator with low diffusivity and strong autocatalytic properties that amplifies initial asymmetries, and a lateral inhibitor, and a more diffusible antagonist that provides negative feedback to confine the pattern to specific regions. This general framework is ubiquitous in biological systems. Canonical examples of excitability also take the form of activator-inhibitor systems [27, 28] where a single equilibrium is stable. Numerous models have been developed to describe the cytoskeletal dynamics [29–31] and signaling behavior in migrating cells [32–37]. Some of these models address the coupled interaction between signaling and the cytoskeleton [38–41], while others focus on how these excitable patterns drive cell movement [38, 42–52]. Previous models have shown that the combination of local positive and global negative feedback can polarize cells [38, 45]. Experiments have suggested that global inhibition could be effected through increased cortical/membrane tension [53–55]. More recently, findings in *Dictyostelium* cells suggest the presence of a local inhibitory feedback loop mediated by actomyosin at the rear of the cell [24, 56]. Our aim is to understand the relative roles of positive and negative feedback loops from the cytoskeleton to the signaling network and how their spatial distribution defines stable patterns within the excitable system.

**Fig. 1.**
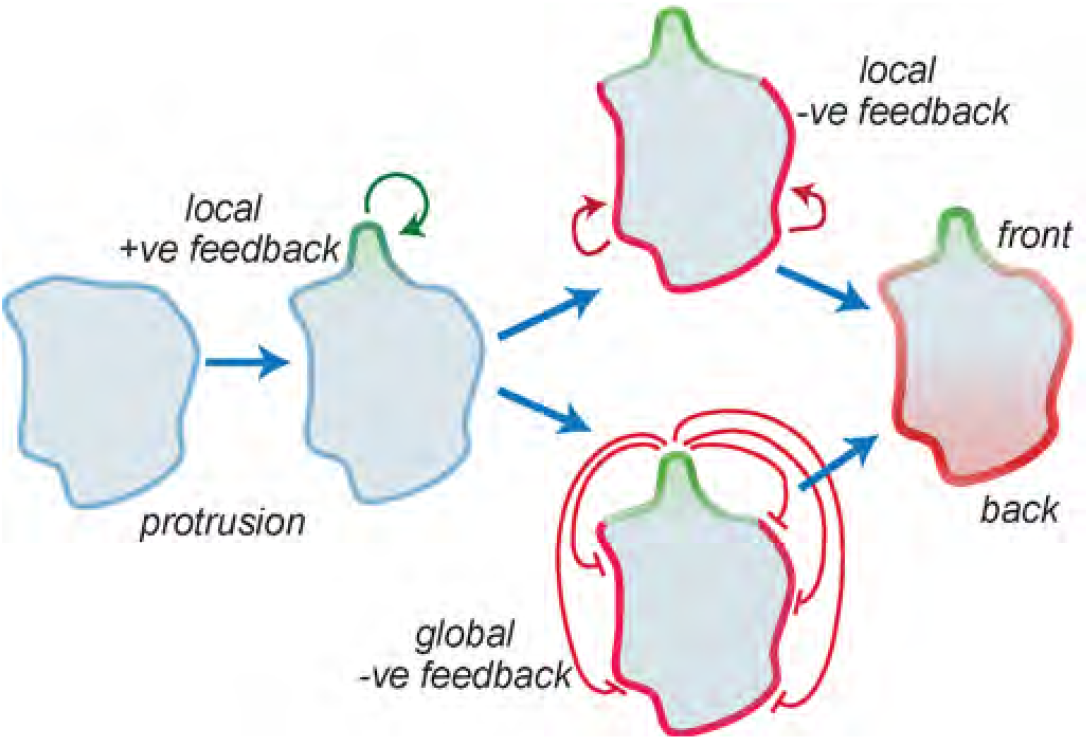
Cytoskeletal-mediated feedback loops influence STEN activity. Regions of elevated Ras lead to localized actin polymerization, initiating a positive feedback loop that further increases Ras production. Additionally, we consider two negative feedback loops: first, protrusions enhance cortical and membrane tension, which globally inhibits STEN activity throughout the cell (Methods). Second, Ras formation is inhibited by the back components and the acto-myosin network, representing a positive feedback loop for rear signaling. The interplay between these feedback loops contributes to the stabilization of the front-rear axis in cells.

## RESULTS

### The core excitable system

To explore cell patterning in the absence of an intact cytoskeleton, we revisited a computational model of the signal transduction system in *Dictyostelium* (Methods). This model (Fig. 2A) consists of three interacting species—Ras, PI(4,5)P_2_, and PKB—that form the core of an excitable network. Ras and PI(4,5)P_2_ exhibit mutually inhibitory interactions, partitioning the membrane into complementary domains: regions of high Ras activity correspond to low PI(4,5)P_2_ levels, and vice versa [57]. This double-negative feedback loop generates the classical positive feedback essential for excitability.

**Fig. 2.**
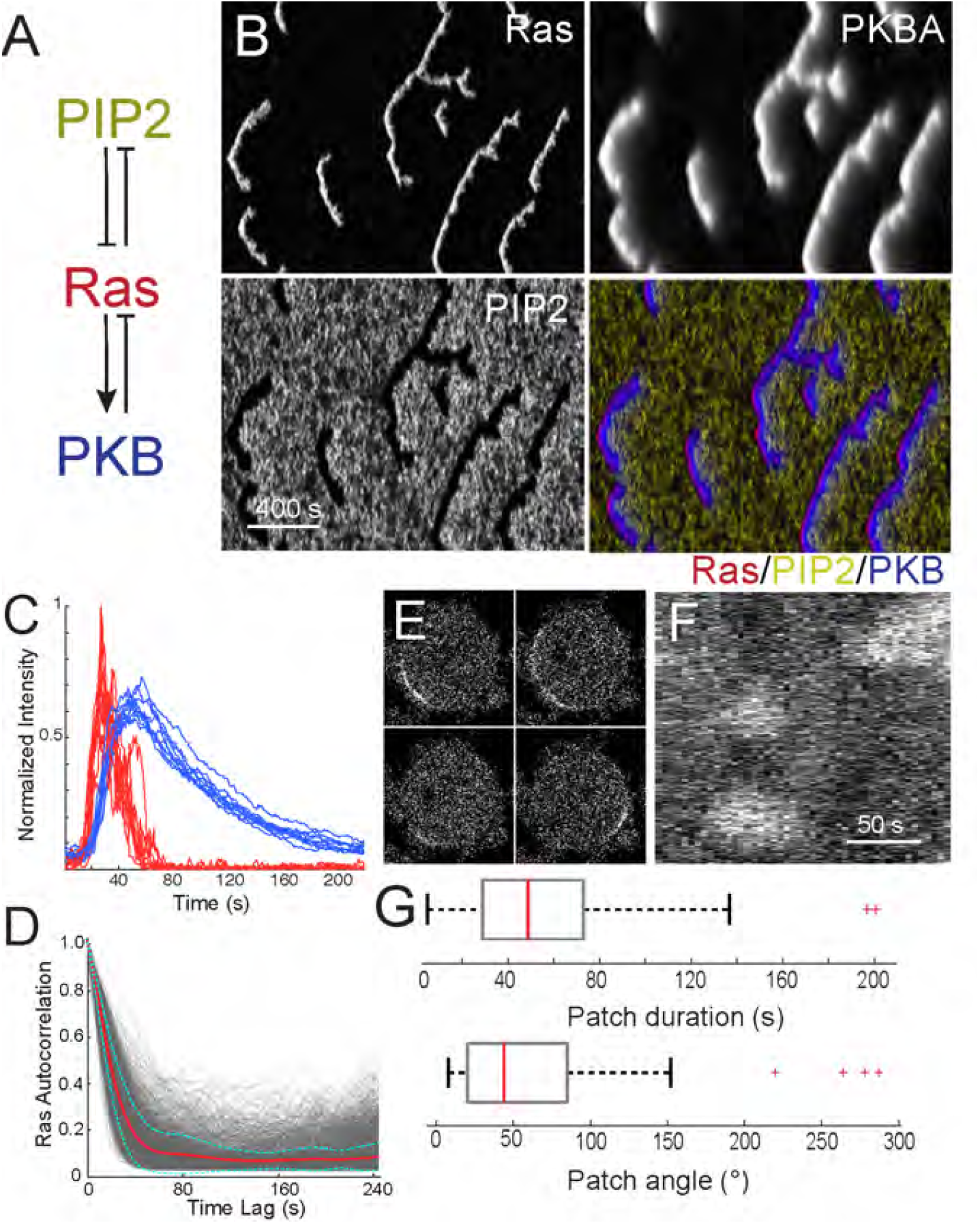
Cells lacking a cytoskeleton display patches with low temporal correlation. A. Schematic of the STEN moodel incorporating Ras, PIP2 and PKB (Methods). B. Representative kymographs of each of STEN components showing spatial and temporal behavior for an unstimulated cell. C. Traces of 16 simulated firings, showing Ras (red) and PKB (blue). D. Ras temporal correlation from simulations (4 simulations; 126 traces each). The red line shows the average of all the traces; the cyan lines are mean ± std. E. Images of a Latrunculin-treated *Dictyostelium* cell displaying RBD-GFP. F. Corresponding kymograph around the perimeter. G. Box plot for the patch duration and angular size for the RBD-GFP patches (*n* = 81 patches in 15 cells).

PKB serves as the network’s refractory component, activated by Ras but providing slow, negative feedback to suppress Ras activity [16]. The model was implemented using coupled stochastic reaction-diffusion equations in both one- and two-dimensional environments (Methods).

Simulations of this model in a one-dimensional periodic domain exhibited occasional firings characteristic of excitable systems. Following these firings, distinct traveling waves emerged from the site of excitation, propagating laterally in both directions and forming V-shaped patterns in kymographs, where the tip of the “V” indicates the initiation point of the wave (Fig. 2B). The stochastic nature of the model resulted in spatial asymmetries, causing one arm of the “V” to extinguish before the other. As depicted in the kymograph, PI(4,5)P_2_ and Ras levels were complementary, with PKB exhibiting patterns similar to those of Ras but with a slower decay. The durations of Ras and PKB firings were 51.6 ± 48.4 s and 96.9 ± 39.8 s, respectively (Fig. 2C; mean ± s.d.; *n* = 159 firings).

In the model, the initiation of a firing is a purely stochastic event, where molecular noise drives the system sufficiently far from the unique dynamic stable equilibrium, resulting in a stereotypical all-or-nothing response typical of excitable systems. To quantify the duration of this response, we measured the temporal autocorrelation of Ras activity at each spatial point in the kymograph (Fig. 2D). This analysis revealed a small correlation, which dropped to 0.1 within approximately 60 s (Fig. 2D). At longer time scales, there was little memory of the initial excitation.

To compare with experimental data, we imaged latrunculin-treated *Dictyostelium* cells and created kymographs of Ras-GFP around the perimeter (Fig. 2E-G). These cells also displayed spontaneous patches of activity around the perimeter, with sizes of approximately 50° and durations of about 60 s.

### Complimentary local feedback loops from the cytoskeleton enhance cell polarity

To determine how feedback loops from the cytoskeleton affect cell polarity, we first incorporated two complementary local feedback loops into our model (Fig. 3A; Methods). We included a positive feedback connection between PKB and Ras, noting that elevated PKB leads to increased actin polymerization, which in turn elevates Ras levels [16]. This positive feedback depends on the amount of branched actin in the cell [57] and increases the likelihood that STEN will fire in the vicinity of a protrusion. Because this process depends on the actin cytoskeleton, we restricted its diffusion so that it acts primarily as a local process. Similarly, we added inhibitory feedback between PI(4,5)P_2_ and Ras to capture the fact that myosin inhibits Ras activation [57]. This was also incorporated as a local loop (Methods).

**Fig. 3.**
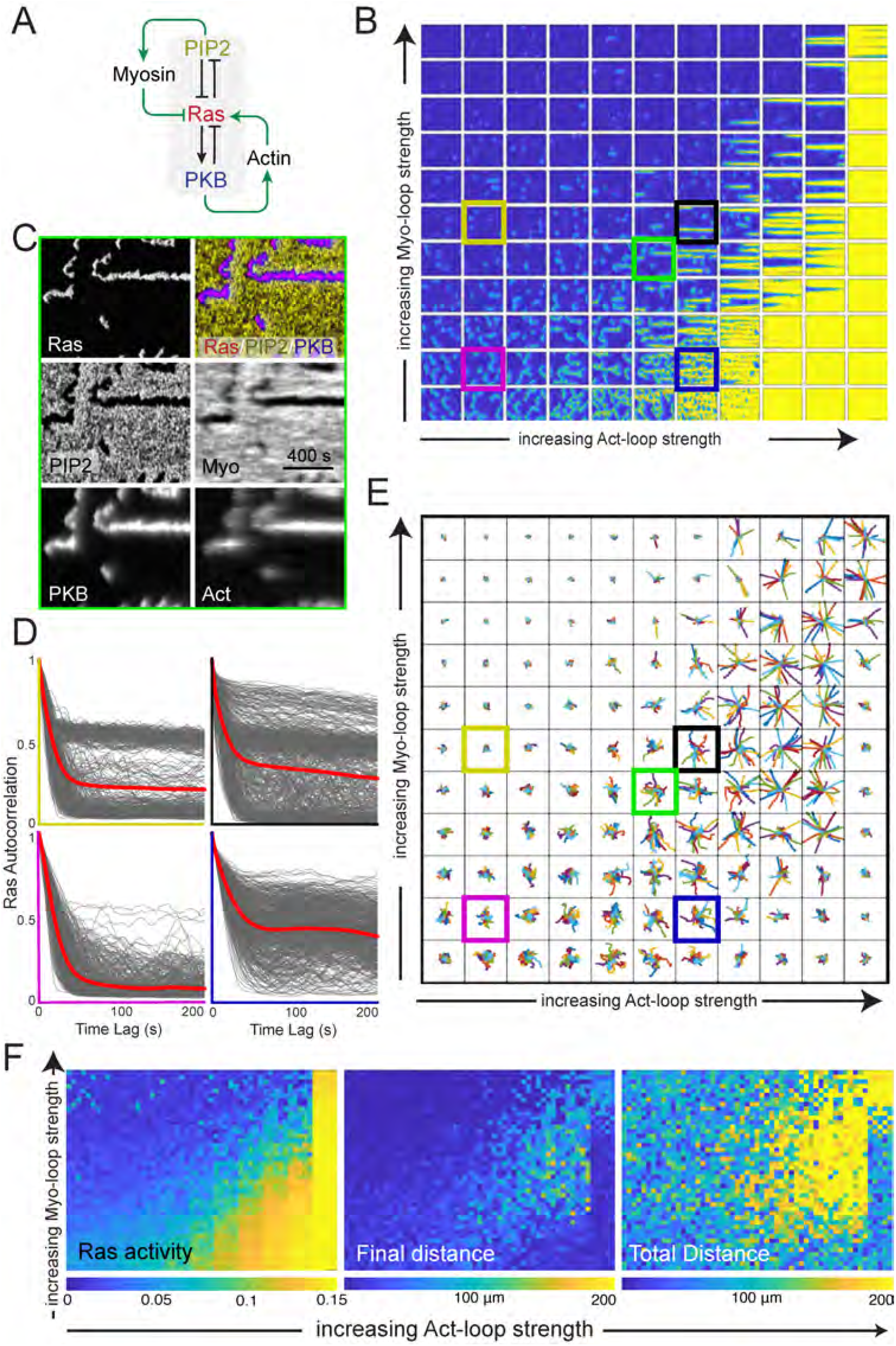
Balancing local positive feedback loops at the front and rear polarize the cell. A. Schematic of the system. The STEN system Fig. 2 forms the core. To this, two positive feedback loops are added which primarily work at the front (Ras to PKB to branched actin back to Ras) and rear (negative from Ras to PI(4,5)P_2_ to myosin and then negative to Ras.) B. Sample kymographs of the system under varying feedback loop strengths. On the horizontal, the strength of the acting loop increases from left to right from 0 to 2 at 0.2 steps. On the vertical, there is a similar increase in the myosin feedback strength, starting at the bottom. Each kymograph shows the activity over 1200 seconds. C. Kymographs of the individual components. This corresponds to the kymograph marked in green in panel B. D. Temporal correlations for four kymographs from panel B. They are colored (yellow, black, magenta and blue) to match those in panel B. E. Trajectories for systems under the varying feedback strengths. Each condition includes 20 simulations. The strengths are as in panel B. F. Heatmaps showing Ras activity (on a logarithmic scale), total distance traveled, and final displacement for the 2420 simulations (20 simulations for each of the 121 different combinations of the feedback strengths. Each condition is plotted as a 4 by 5 matrix.

To investigate how cytoskeletal feedback loops influence cell polarity, we incorporated two complementary local feedback loops into our model (Fig. 3A). First, we introduced a positive feedback loop between PKB and Ras, based on the observation that increased PKB promotes actin polymerization, which in turn enhances Ras activity [16]. This feedback depends on the abundance of branched actin [24] and increases the likelihood that STEN will fire near a protrusion. Since this mechanism relies on the actin cytoskeleton, we restricted its diffusion, ensuring it functions as a local process. Similarly, we added an inhibitory feedback loop between PI(4,5)P_2_ and Ras to account for myosin-mediated suppression of Ras activation [57], implementing this loop locally as well (Methods).

To assess the relative contributions of these feedback loops, we systematically varied their strengths, simulated the system, and visualized the results as a kymograph matrix. In this matrix, the strength of the branched actin feedback loop increased along the horizontal axis, while the myosin-mediated inhibition increased along the vertical axis (Fig. 3B). As previously observed, some simulations produced kymographs where waves translocated laterally following an initial stochastic trigger (e.g., kymograph in magenta). For a fixed myosin feedback strength, increasing actin feedback led to heightened Ras activity along the cell perimeter, generating more and longer-lived patches. Conversely, at a constant actin feedback strength, increasing myosin feedback resulted in reduced activity.

Notably, when both feedback loops reached moderate strengths (e.g., kymograph marked by the black box), we observed stable, persistent patches. In this regime, once the system fired, activity remained elevated without spatial translocation along the membrane. At high positive feedback levels, the entire cell boundary became uniformly active, sustaining elevated activity throughout the simulation, even under strong inhibitory feedback. For moderate positive feedback strengths, increasing the strength of the actomyosin negative feedback loop lowered activity and broke persistent patches into lesser durable patches, reducing polarization.

Kymographs of individual model components (Fig. 3C) revealed that actin and myosin feedback loops aligned with the front (Ras) and rear (PI(4,5)P_2_) signaling modules, respectively, demonstrating that both loops reinforced polarity in complementary membrane regions.

Plots of the temporal Ras autocorrelation for four different combinations of feedback loop strengths demonstrated that increasing either feedback loop strength led to more correlated behavior (Fig. 3D). At high actin feedback levels (blue kymograph), the correlation plateaued around 0.35. While stronger myosin feedback resulted in slightly lower values, the overall correlation remained elevated.

To assess the impact of these feedback loops on cell motility, we used the kymographs (20 per feedback strength combination) to simulate cell movement (Fig. 3E; Methods). The results revealed complex biphasic behavior. In the left region of the matrix, where front-feedback strength was low, increasing negative feedback suppressed motion. Conversely, along the bottom of the matrix, where rear feedback was weak, increasing positive feedback enhanced motility. However, in cases where kymographs displayed uniformly high Ras levels (last column), cell motility was inhibited. In these conditions, increasing rear inhibition facilitated efficient migration.

To quantify these effects, we generated heatmaps for total Ras activity, total distance traveled, and net displacement across all simulations (Fig. 3F). The heatmaps revealed that increasing actin or myosin feedback led to higher or lower Ras activity, respectively. This trend largely corresponded to total distance traveled, except at extreme Ras activation levels, where high positive feedback and weak negative feedback impaired motility. Finally, the highest net displacement occurred when both feedback loop strengths were elevated.

### Local positive and global negative feedback loops reduce the presence of multipolarity

We next explored a model in which actin is involved in complementary positive and negative feedback loops (Fig. 4A). The motivation for the positive feedback was consistent with previous findings: increased levels of Ras and PKB lead to greater actin polymerization, which in turn further elevates Ras levels. For the negative feedback, we were inspired by reports indicating that actin polymerization increases membrane and cortical tension, which inhibits signaling at the cell front [54, 55].

**Fig. 4.**
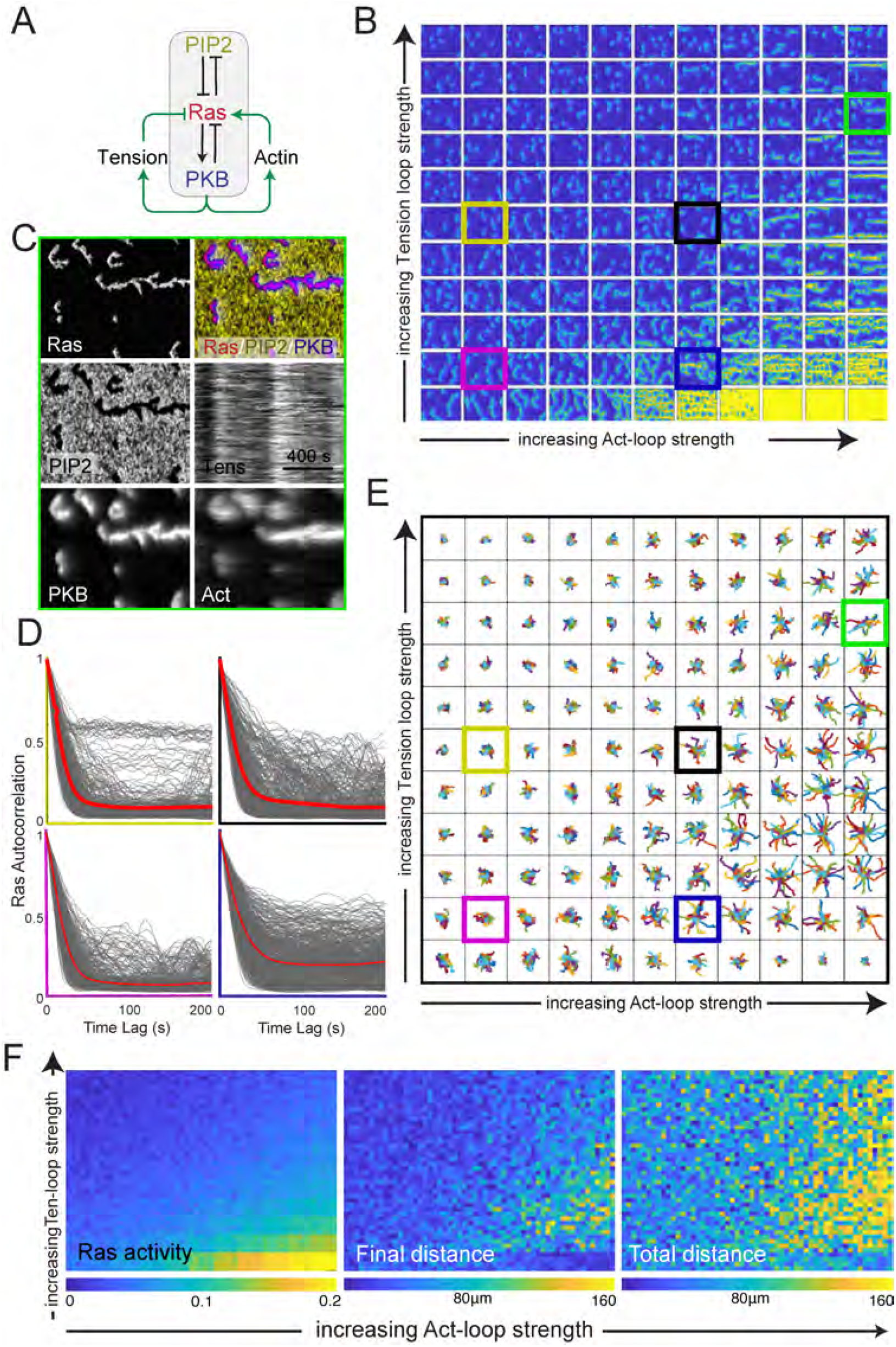
Global inhibition can reduce multipolarity. A. Schematic of the system. The STEN system Fig. 2 forms the core. To this, two feedback loops are added that come from protrusions. The local enhancement is as in Fig. 3. Additionally, we include a negative feedback loop representative of cortical/membrane tension. B. Sample kymographs of the system under varying feedback loop strengths. The panel is as in Fig. 3 except that the negative feedback strength is from 0 to 4.0 with step size of 0.4. C. Kymographs of the individual components. This corresponds to the kymograph marked in green in panel B. D. Temporal correlations for four kymographs from panel B. They are color-cide (yellow, black, magenta and blue to match those in panel B. E. Trajectories for systems under the varying feedback strengths. Each condition includes 20 simulations. The strengths are as in panel B. F. Heatmaps showing Ras activity (logarithmic scale), total distance traveled, and final displacement for the 2420 simulations (20 simulations for each of the 121 different combinations of the feedback strengths. Each condition is plotted as a 4 by 5 matrix.

Kymographs of the system under varying feedback loop strengths (Fig. 4B) revealed similar patterns to our previous model: increasing local positive feedback resulted in more and longer-lasting firings, while increasing negative feedback suppressed this response. The optimal polarity—characterized by single, longlived streaks—occurred when both feedback levels were high (e.g., the kymograph marked in red). Kymographs of the individual components showed that actinmediated feedback correlated both temporally and spatially with PKB, whereas the tension-mediated negative feedback exhibited temporal correlation with PKB but swept across the cell spatially (Fig. 4C).

The autocorrelation patterns in these simulations were similar to those observed in the earlier model, though high-tension feedback induced more pronounced bistability, with clearer distinctions between high and low correlated states compared to the previous model (c.f. Fig. 4D vs. Fig. 3D, particularly in the kymographs marked by the yellow and red boxes). The impact of the feedback loops on cell motility (Fig. 4E, F) mirrored patterns from the earlier model. We observed a similar biphasic response in scenarios with high positive feedback and low negative feedback, but this effect diminished with minimal negative feedback.

### Morphological differences drive cell motility efficiency

To examine the correlation between cell shape (polarity) and cell movement, we imaged unstimulated *Dictyostelium* cells and quantified their circularity. We identified three broad classifications: cells exhibiting long, narrow shapes (circularity values less than 0.25; Fig. 5A; S1 Video), cells with little eccentricity (circularity values greater than 0.45; Fig. 5C; S3 Video) and an intermediate group of cells, occasionally displaying some polarized morphology, with circularity values between these

**Fig. 5.**
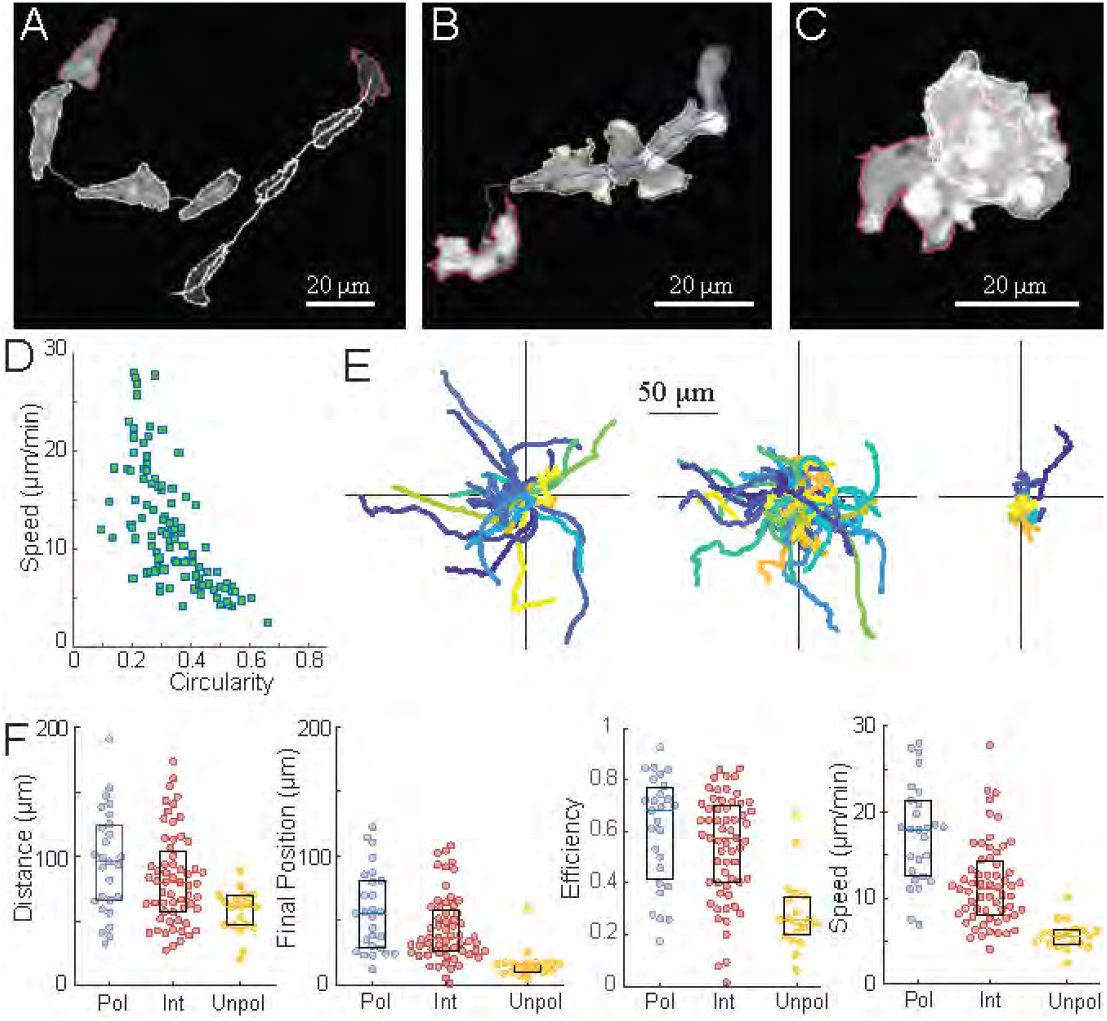
Correlation between shape and movement. A-C. *Dictyostelium* cells were imaged over time, and their shape and displacement were measured over time. Cells displayed polarized (A; S1 Video), intermediate (B; S2 Video) or unpolarized morphologies (C; S3 Video). The snapshots here show cells 120 seconds apart. D. Correlation between speed and circularity. E. Displacement over time for cells in the three classes. These were chosen based on the average circularity over time. Polarized cells have circularity less the 0.25; intermediate cells have circularity between 0.25 and 0.45; and unpolarized cells have circularities greater than this. F. Total distance, final position, efficiency and speed for the various cells in the three classes. Total number of cells were 27, 60 and 19, respectively.

0.25 and 0.45; Fig. 5B; S2 Video). To correlate these cellular morphologies with cell movement, we computed the total distance traveled, final position, efficiency (the ratio of the previous two values), and speed of the cells and found the correlation with circularity (Fig. 5D). The corresponding Pearson correlations were *ρ* = −0.34, *p* < 3 × 10−4 (total distance traveled), *ρ* = −0.48, *p* < 2 × 10−7 (final position), *ρ* = − 0.51, *p* < 2 × 10−8 (efficiency) and *ρ* = − 0.67, *p* < 3 × 10−15 (speed) from *n* = 106 cells. Polarized cells moved in relatively straight lines (Fig. 5E) and traveled the farthest distances (Fig. 5F). When these cells turned, they did so in gradual arcs. At the opposite extreme, cells with little eccentricity despite being active and displaying deviations from a circular shape, did not maintain a consistent form. In this case, the centroid did not exhibit significant movement. Finally, cells in the intermediate group moved a moderate distance, but they often came to a stop. Occasionally, they would adopt a shape perpendicular to their previous movement.

### Polarization potential is compromised for local inhibition model

The mechanisms in Fig. 3 and Fig. 4 can both produce polarized kymographs and sustain persistent, directional migration, provided that the positive and negative feedback loops are properly balanced. To investigate this further, we analyzed several polarization markers (Fig. 6).

**Fig. 6.**
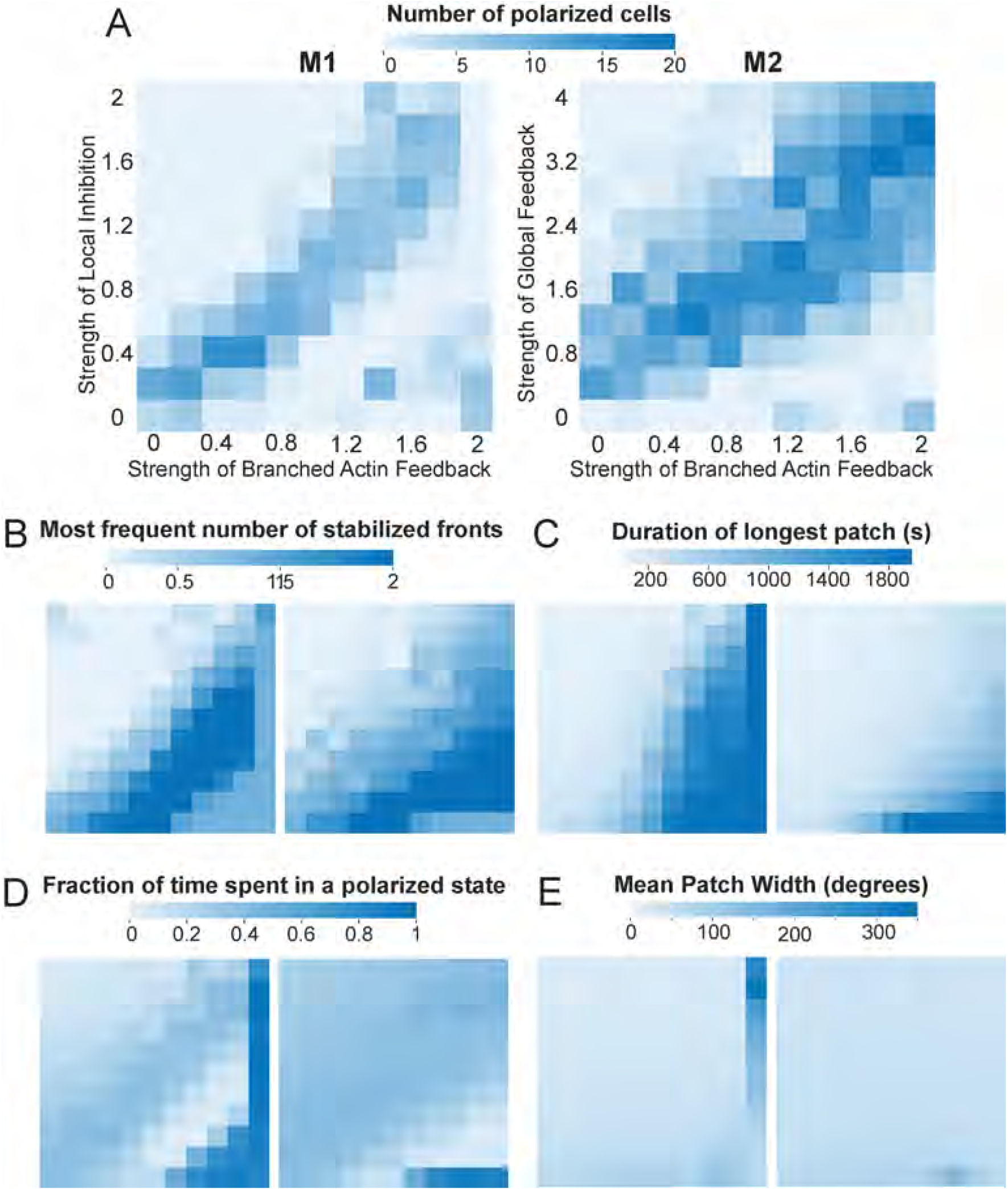
Heatmaps showing polarization potential of the two models. The left column shows statistics for Model 1 (M1; local inhibition), and the right for Model 2 (M2; global inhibition). The *x*-axis for both M1 and M2 shows strength of Branched Actin (BA) feedback loop ranging from 0 to 2.0, in 0.2 intervals. For M1, the *y*-axis is the strength of myosin feedback ranging from 0 to 2 in intervals of 0.2; for M2, it is the strength of the global inhibition loop, ranging from 0 to 4 in intervals of 0.4. The difference in these ranges accounts for differences in inhibitor levels in the two systems such that strength of negative feedback on STEN stayed the same. A. Number of polarized cells in a population size of *n* = 20 cells. In each simulation, patches are filtered to include only those that have more than 50 pixels and the ellipse fit to the patch made an angle of less than 30° with respect to the time axes.At each time step, the number of filtered patches was quantified. B. The most frequently appearing number of patches at any point in time. C. Length of the longest patch, in seconds. D. Fraction of time spent in a polarized state. This uses the data from panel B to obtain the time points for which there is a single front. The sorting criteria were based on outcomes from panels B–D, where “polarized” cells exhibited a single front as their most frequent state and the mean patch width was less than 60^°^. E. Mean path width, in degrees.

For example, in Fig. 6A, we compared the fraction of cells exhibiting polarized kymographs. Cells modeled using Model 1 (local inhibition; Fig. 3) displayed significantly less polarization compared to Model 2 (global inhibition; Fig. 4). In both models, polarization predominantly occurred along the diagonal in the branched actin and myosin strength grid. In Model 1, the highest number of polarized cells (15–16) was observed at branched actin strengths of 0.4–0.6 and myosin strengths as low as 0.4, after which polarization declined to 10 or fewer cells. In contrast, Model 2 exhibited a broader range of polarization, with counts between 15 and 20 along the diagonal. Additionally, in Model 2, the polarization region expanded toward the top right corner, suggesting that increasing branched actin strength and global inhibition enhances polarization.

We further compared the models by analyzing the most frequent number of stabilized fronts in each kymograph (Fig. 6B). Model 1 produced several instances of the “2-front cell” along the diagonal of increasing branched actin and myosin strength. In contrast, Model 2 exhibited most of its 2-front states in the high branched actin, low global inhibition zone. Notably, achieving a “single front” cell for a given level of positive feedback required higher feedback strengths in Model 1 compared to Model 2. This distinction is reflected in the polarization potential heatmap (Fig. 6A), where only cells that predominantly exhibited a single front are included.

Examining the duration of polarity patches, we found a clear positive correlation between the longest persisting patch and the strength of branched actin feedback (Fig. 6C). Notably, Model 1 exhibits longer patch durations when branched actin strengths exceed 1.0 compared to Model 2.

To quantify the fraction of time spent in a polarized state, we computed the proportion of simulation time during which cells maintained a single front (Fig. 6D). In Model 1, cells remain polarized for approximately 40– 60% of the simulation time along the diagonal, indicating increased polarization when feedback strengths are balanced. In Model 2, this effect is less pronounced, as polarization occurs across a broader range of feedback strengths. Extremely high polarization values (≥ 80%) correspond to conditions in Fig. 3 and Fig. 4, where Ras activity stabilizes at a persistently high level. In these cases, cells exhibit a single high-activity patch that spans the entire membrane, resulting in the longest patch durations (Fig. 6C). However, these cells are not polarized in the traditional sense and are unable to migrate (Fig. 3, Fig. 4).

### Local, but not global inhibition captures the behavior of cells under perturbation

While both local and global inhibition models generate polarization across different parameter spaces, we further examined their biological relevance by testing their ability to replicate perturbation experiments. These experiments involve overexpression or underexpression of key molecules in the signaling pathway, as well as optogenetic modulation through laser-induced recruitment or depletion of proteins in wild-type cells. Perturbations targeting different polarization loops have been extensively studied in *Dictyostelium* [24]. Here, we analyze similar perturbations in neutrophils, which exhibit analogous responses, emphasizing the universality of polarization mechanisms.

A wild-type polarized neutrophil is characterized by a flared-out front and a narrow tail at the back. Actin fluorescence is observed at both ends: branched actin at the front and actomyosin-rich, contractile structures at the rear.

Perturbations are introduced at time zero, with cellular responses tracked over time. In the first experiment, Ras activity was globally increased by activating a 488 nm laser to recruit cytosolic KRas4B G12V to the cell periphery (Fig. 7A). Before recruitment, both cells exhibited single, stable fronts and directional movement (S4 Video). In the second frame, the top cell changed direction but retained its front. Following Ras recruitment, multiple sites along the membrane became active upon cell-cell contact, ultimately leading to a two-front phenotype in the top cell. In silico, Ras upregulation produced different outcomes depending on the polarization model. In the local excitation–local inhibition model, increased Ras induced widespread, persistent activity across the membrane, disrupting polarity. In contrast, in the local excitation–global inhibition model, Ras enhancement reinforced localization of a single persistent patch, promoting stronger polarization.

**Fig. 7.**
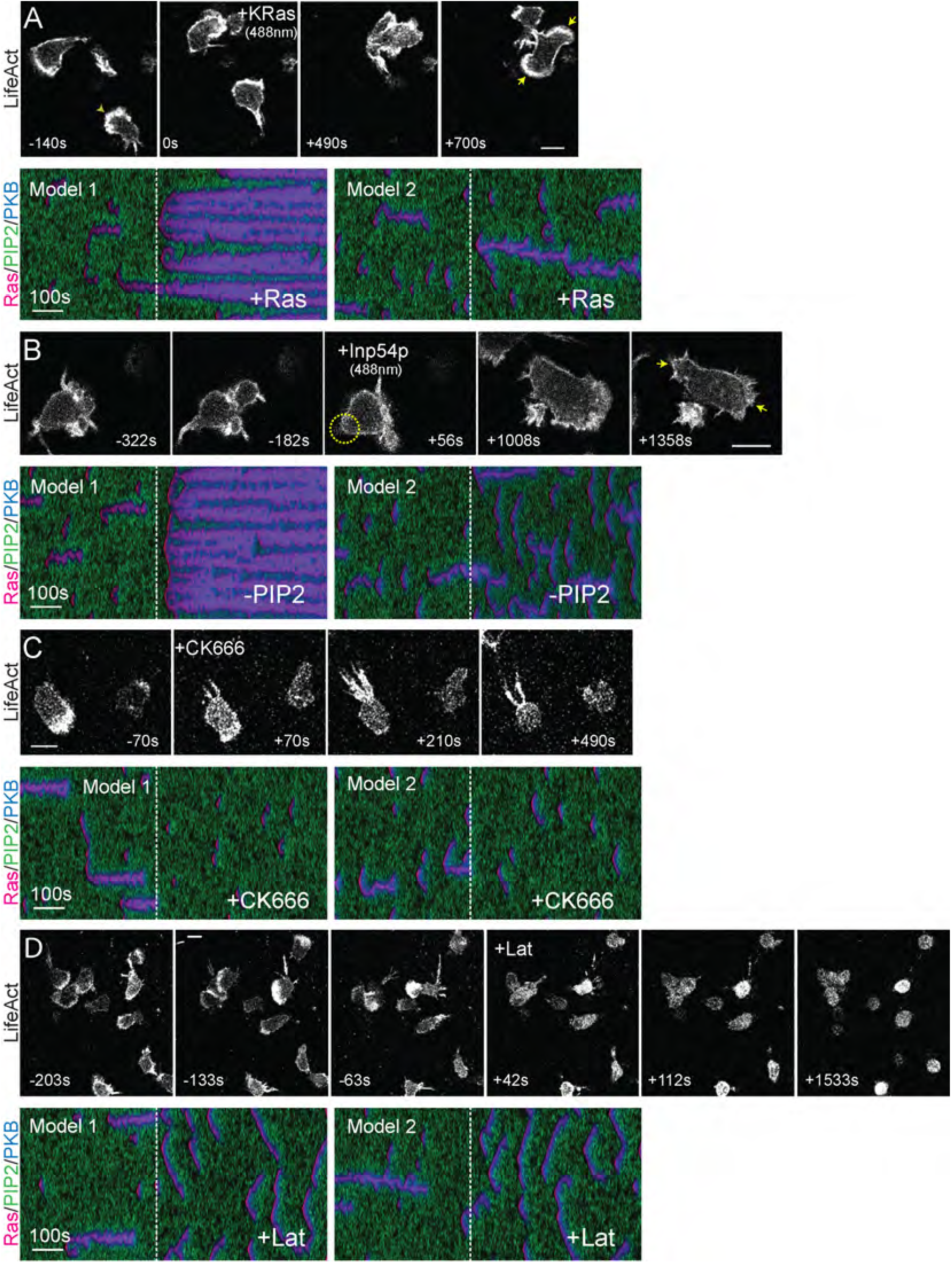
Perturbations that impact front and back polarity. (A-D) Differentiated neutrophils with F-actin-rich fronts (denoted by LifeAct-miRFP703) were imaged over time, and their shape and displacement were measured. Corresponding perturbations were simulated using Model 1 (local inhibition) and Model 2 (global inhibition), with the first 400 seconds showing STEN activity before effector recruitment (A–B) or drug addition (C–D), and the last 500 seconds showing STEN activity post-effector recruitment (A– B) or drug addition (C–D). The three colors in the kymograph represent Ras (Red), PKB (Blue), and PIP2 (Green). A. A 488 nm laser was switched on at frame 50 to globally recruit cytosolic CRY2PHR-mCherry-KRas4B G12V ΔCAAX to the cell periphery, increasing Ras activity and activating the STEN network. B. A 488 nm laser was switched on at frame 47 to globally recruit cytosolic CRY2PHR-mCherry-INP54P to the cell periphery, reducing PIP2 levels, which serves as the back molecule in STEN in both models. C. Differentiated neutrophils were pre-treated with 50 *µ*M CK666 to inhibit branched actin (denoted by LifeAct-miRFP703), turning off its positive feedback. D. Cells were pre-treated with 5 *µ*M Latrunculin B to completely remove actin polymerization (denoted by LifeAct-miRFP703), turning off both positive and negative feedback. Scale bar: 5 *µ*m.

Next, we tested the effects of PIP2 depletion by globally recruiting cytosolic INP54P to the membrane using a 488 nm laser (Fig. 7B, S5 Video). Following recruitment, the actomyosin tail disappeared, and the leading edge expanded. The cell developed a second front at its former rear, with no reformation of the actomyosin tail. In silico, PIP2 depletion had distinct effects in each model. In the local excitation–local inhibition framework, its impact resembled that of Ras upregulation, leading to loss of polarity and widespread activity. In the local excitation–global inhibition model, polarization decreased due to the formation of broader, less persistent, and more dispersed waves.

The first two perturbations targeted components of the signal transduction excitable network (STEN). In the third experiment, we inhibited Arp2/3-mediated branched actin formation using CK666 (Fig. 7C, S6 Video), directly disrupting the positive feedback loop between the cytoskeleton and STEN. Following inhibition, the leading edge collapsed, while the actomyosin-rich rear enlarged and intensified in fluorescence. Since actin could no longer form branched networks, excess monomeric actin accumulated at the rear, reinforcing actomyosin structures. The resulting loss of cytoskeletal feedback disrupted front persistence and patch localization in both models, severely impairing motility.

Finally, to abolish all cytoskeletal feedback, we treated cells with Latrunculin B, an inhibitor of actin polymerization (Fig. 7D, S7 Video). Wild-type cells with well-defined fronts and tails began to round up upon drug exposure. The actomyosin tail and leading edge retracted, and after prolonged treatment, cells became entirely spherical. In silico, removal of both positive and negative feedback loops caused both models to revert to the basal STEN phenotype observed in Fig. 2, underscoring that the observed differences arise solely from cytoskeletal feedback interactions.

### Dynamic partitioning can generate local and global inhibition

To explore the mechanisms underlying cell polarization further, we next developed a model based on *dynamic partitioning*, the principle that signaling protein distributions are dynamically regulated to create distinct functional domains [58]. Deng *et al*. recently showed that this principle governs the mutually inhibitory localization of PIP5K and Ras [59]. They observed that, in resting cells, PIP5K was uniformly distributed, while Ras activity was minimal. Spontaneous PIP5K displacements disrupted this symmetry, activating Ras and downstream signaling, including PI3K. PIP5K knockout increased Ras-PI3K activation and cortical wave formation, affecting cell protrusion and migration. Conversely, low PIP5K overexpression promoted polarity, highlighting its role as a key regulator. We incorporated these findings into our model.

We then examined PIP5K’s inhibitory role and its potential contribution to myosin-mediated local inhibition (Fig. 8A). Total PIP5K, consisting of bound and unbound states, was conserved. Both states contributed to PI(4,5)P_2_production, inhibiting Ras.

**Fig. 8.**
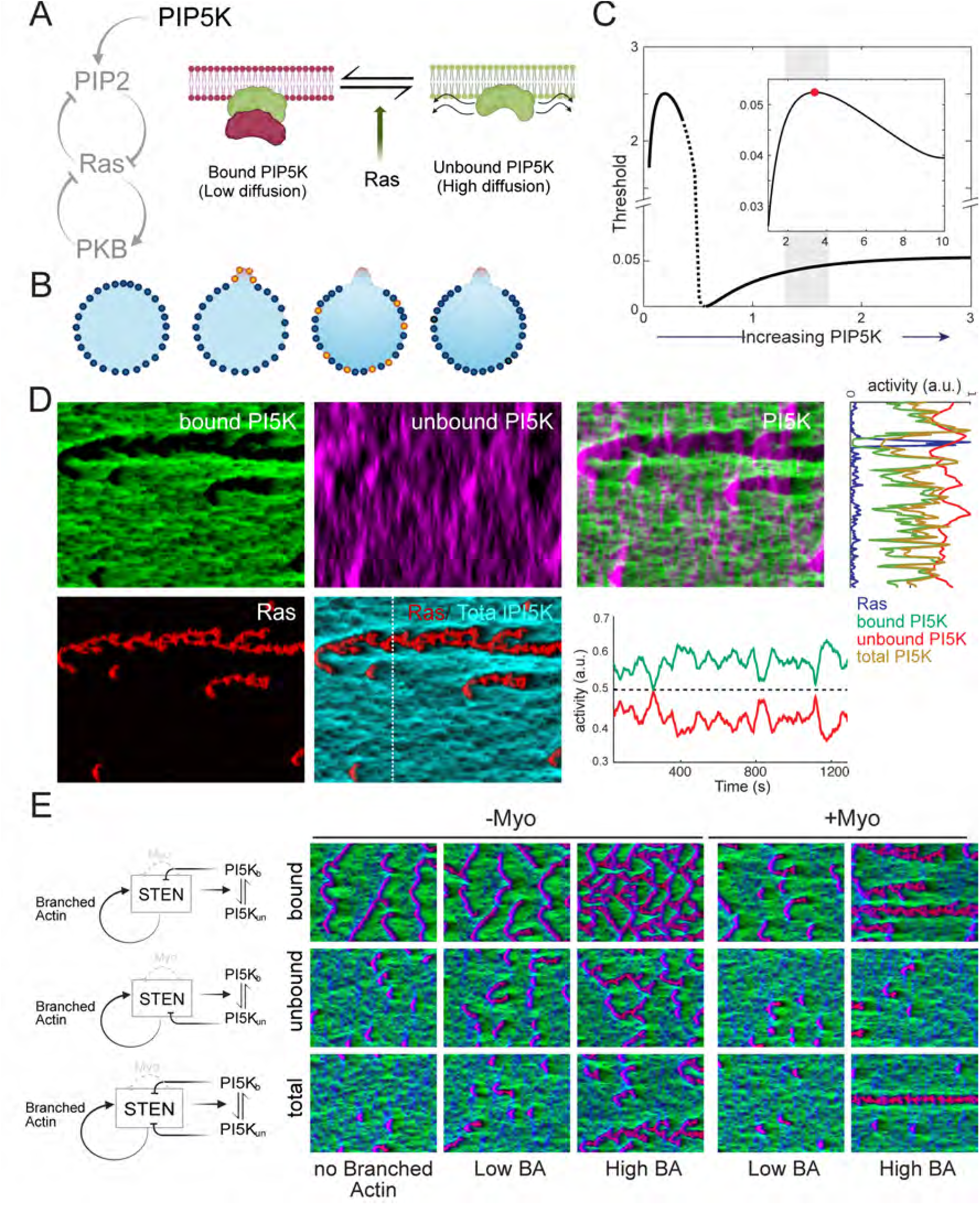
Dynamic partitioning mechanism of PIP5K can increase polarization potential. A. Scheme of PIP5K action on the STEN excitable network. B. Stages of the PIP5K action during protrusion formation. The cell starts with having all bound PIP5K (blue dots). When a protrusion is formed front markers unbind PIP5K (orange0, which displaces to the back due to high diffusion, where it converts back to the bound state. C. Effect of PIP5K levels on the threshold. The inset shows a zoomed-in view of the threshold for PIP5K levels ranging from 1 to 10. A value of 1 corresponds to the basal state, without additional recruitment of PIP5K. The shaded region (1.6–1.7) represents the elevated PIP5K levels explored in our simulations. An increase from the basal value of 1 to any value within this range results in a higher threshold, as indicated by the positive slope of the curve. D. Simulation of PIP5K action in a cell with polarity. Simulation type: Total PIP5K inhibition, BA strength 1.0, myosin strength 0.4. E. Schematics and simulations illustrating the effects of different PIP5K contributors to inhibition: inhibition via bound PIP5K (**Bound**), unbound PIP5K (**Unbound**), and an equal contribution from both forms (**Combined**). In the simulation grid, rows correspond to BA strengths: 0 (No BA), 0.2 (Low BA), and 0.8 (High BA); columns correspond to myosin strengths: 0 (-Myo) and 0.4 (+Myo). This arrangement applies to all simulations in the grid except for the bottom-right corner (High BA, +Myo), where the highly polarized phenotype corresponds to BA strength 1 and myosin strength 0.6.

In simulations of unperturbed cells, PIP5K was primarily in the bound state. Protrusion formation triggered Ras-mediated conversion of PIP5K to the unbound state, which diffused rapidly owing to its higher diffusion coefficient (Fig. 8B). The unbound form dissipated from the front and, through reversible conversion (Fig. 8A), returned to the bound state.

Our model posits that PIP5K increased PI(4,5)P_2_ production. Varying PI(4,5)P_2_ levels revealed its impact on the activation threshold. A value of one corresponded to the absence of PIP5K, with the gray region in Fig. 8C indicating its effect. PIP5K increased the production rate from 1.4 to 1.7, as confirmed by simulations (Fig. 8C). The inset further validated the elevated threshold (production rates 1-10). Thus, PIP5K raised the system’s activation threshold.

The two PIP5K states exhibited distinct distributions (Fig. 8D). Bound PIP5K was depleted at the Ras-active front but enriched at the periphery and rear, with limited spatial spread owing to low diffusion. Unbound PIP5K appeared at the front and dissipated rapidly. Increased bound PIP5K regions coincided with the termination of unbound PIP5K waves. Bound and unbound PIP5K concentrations remained complementary, maintaining a constant total concentration of one. The time course (dotted line) confirmed that half of the total PIP5K was conserved. The spatial profile across the white dotted line in the kymograph showed a dip in bound PIP5K at Ras firing, consistent with the kymograph.

Finally, we examined PIP5K inhibition mechanisms (Fig. 8E). Bound PIP5K had minimal effect on wave patterns, regardless of myosin or branched actin levels. In contrast, unbound PIP5K exerted a uniform inhibitory effect. Persistent patches emerged only under high branched actin levels and absent myosin inhibition. When both PIP5K states contributed equally to inhibition, persistence emerged. Even at low levels, a memory effect was observed, with waves recurring at the same location. Branched actin positive feedback amplified this effect. At high branched actin levels, a primary and secondary persistent streak appeared. Myosin suppressed activity at regions of low branched actin, preventing polarity. However, with high branched actin levels, a single dominant streak emerged, indicating clear polarization.

## DISCUSSION

In one of the earliest computational studies on migratory cell polarization [60], Meinhardt proposed that three processes were required. The first two mirror the classical Turing/Meinhardt-Gierer model of pattern formation: (1) a local self-enhancing reaction and (2) a longrange (global) inhibitor that counterbalances it. Spatial localization arises when activation occurs at the membrane or cortex, while inhibition diffuses freely across the membrane. Many early models of chemoattractant-induced polarization were based on these two feedback loops [61, 62], which together generate diffusion-driven stable patterns. In the context of cell motility, these localized patches of high activity give rise to persistent, directional motion. However, in the absence of an external guiding gradient, cells do not typically remain polarized. To address this, Meinhardt introduced a third component: a local, longer-lived antagonist of the activator that destabilizes high-activity regions, allowing new ones to form. Since Meinhardt’s original work, substantial experimental evidence has shown that cell motility is achieved through firings of the cytoskeletal excitable network and that this process is directed by an independent signaling network. However, although a model combining local positive feedback with local and global negative feedback seems plausible and leads to realistic cellular motion [45], there is little experimental evidence to support it.

Positive feedback from the cytoskeleton to the signaling network plays a crucial role in establishing polarization by amplifying signaling activity and creating a spatial memory of prior excitations, thereby increasing the likelihood that future triggers will activate those regions. This process leads to persistent excitation in specific domains of the cell membrane, which is essential for directional migration. In our models, positive feedback amplifies Ras activity by enhancing its production rate. This positive feedback has a longer lifetime than that of the signaling network, establishing a locational memory following protrusion formation. However, if this memory does not trigger further firings, the effect of the positive feedback eventually decays, causing the memory to fade—*Dictyostelium* cells exhibit a persistence time on the order of several minutes.

Positive feedback alone, however, is insufficient to establish a polarized cell. Cells relying solely on positive feedback often develop multiple stabilized fronts and, in extreme cases, become hyperactive (bottom row, Fig. 3B and Fig. 4B). To counteract this, models have typically incorporated global inhibition to ensure that, once a front forms, the rest of the cell remains quiescent [63–67]. Models based on this local positive, global negative feedback paradigm successfully achieve polarization in both unstimulated cells and those sensing a chemoattractant gradient. In Shi et al. [38], we demonstrated that this mechanism functions within an excitable network (EN) context, guiding movement for cells in gradients and enabling persistent random migration in their absence. The model in Fig. 4 is based on this local positive and global negative feedback paradigm. Experiments suggest that the fast global inhibition required by these models can be achieved through the membrane or cortex. This inhibition typically originates from the strain caused by protrusive forces at the cell’s front, which makes it more difficult for new protrusions to form elsewhere [53–55].

Recent experiments have shown that increasing Ras-GAP in cells with initially high Ras activity enhances polarity [56]. This finding is somewhat puzzling, as Ras-GAP inhibits RasGTP, thus acting as a negative modulator of activity (S1 Fig.). Front-mediated global inhibition fails to account for these observations. More recently, Kuhn et al. identified distinct positive and negative components of cytoskeletal inhibition by perturbing polarized cells [24]. They found that negative feedback originates from actomyosin, establishing a back-mediated inhibition system that localizes at the cell’s rear. Thus, actomyosin enhances polarization by suppressing protrusions at the back.

In our excitable network, which models Ras, PIP2, and PKB as the front, back, and refractory components, respectively, Ras deactivation (RasGTP to RasGDP) is governed by its degradation rate. Thus, GAP directly influences Ras degradation in our equations (Methods). In the model of Fig. 3, local inhibition was used to capture these GAP-related experiments, incorporating “actomyosin” as a local inhibition mechanism originating from PIP2 (back). This mechanism increases Ras degradation more at the rear (where PIP2 levels are high) than at the front (where PIP2 levels are low). In the GAP recruitment experiments, a global increase in GAP led to an overall reduction in Ras activity and more polarized cells [56]. However, in simulations, globally altering Ras degradation in the excitable network failed to enhance polarization unless front-mediated local positive feedback was counterbalanced by actomyosin-driven local negative feedback [56].

While the local inhibition model successfully captured Ras perturbation experiments, it failed to polarize cells robustly—defined as maintaining a single persistent front while suppressing activity elsewhere along the cell perimeter. Even under optimal feedback strengths, the model frequently stabilized two fronts instead of one (Fig. 6). Moreover, compared to global inhibition, local inhibition proved to be a weaker mechanism at higher levels of positive feedback. Whereas global inhibition effectively maintained confined patches of activity and often supported polarization at elevated branched actin levels, local inhibition, at the same mean inhibition strength, became increasingly hyperactive as branched actin strength increased. Eventually, at the highest levels of branched actin feedback, the entire cellular perimeter remained persistently active, resembling conditions observed in the absence of inhibitory feedback (cf. right column vs. bottom row of Fig. 3B). This highlights the critical role of spatial delocalization and long-range negative feedback in establishing stable polarization. To bridge the gap between a model that captures experimental perturbations and one with strong polarization potential, we propose here a scheme based on dynamic partitioning [58]. Recent experiments have shown that PIP5K, a membrane-bound Ras inhibitor, remains confined to the membrane, does not enter the cytosol, and diffuses slowly across the membrane [59].

Based on these experiments, we propose a model for polarity based on PIP5K (Fig. 8). It assumes that PIP5K exists in two states: a slow-diffusing “bound” state (S3 Fig., S8 Video) and a highly diffusible “unbound” state (Fig. 8). The total PIP5K in the system is conserved, but the two states coexist dynamically. High Ras activity at the front causes PIP5K to unbind, allowing it to diffuse away. Binding sites at the rear of the cell capture this unbound PIP5K, enabling it to accumulate there.

At the front, PIP5K unbinds and, in this unbound state, transiently inhibits Ras, interfering with the branched actin positive feedback loop and thereby disrupting persistent patches, which reduces cellular polarization (middle row of Fig. 8E). Therefore, if inhibition is solely accomplished by unbound PIP5K, cells do not polarize. Conversely, if inhibition comes solely from the bound state, which is localized at the rear, it is ineffective due to its highly localized nature and limited diffusion, preventing it from suppressing secondary and tertiary front formations (top right simulation of Fig. 8E). Simulations showed that the most polarized cells were observed when both PIP5K states inhibit Ras (bottom right panel of Fig. 8E). Note that positive feedback from branched actin is required.

We also simulated the system under conditions where myosin is absent so that the only inhibitory mechanism is through PIP5K (-Myo simulations of Fig. 8). These simulations showed that, with positive feedback and both states of PIP5K inhibiting Ras, cells could polarize, but not as efficiently as with the myosin contribution. These simulations recapitulated experimental findings [59]. Thus, we conclude that the combination of branched actin positive feedback, back-mediated myosin inhibition, and PIP5K dynamic partitioning leads to highly polarized cells. Notably, by having both states act as inhibitors, PIP5K achieves both local and global inhibition. When paired with back-mediated myosin inhibition, dynamic partitioning enhances negative feedback at the back, as both PIP5K states inhibit Ras activation while promoting PIP2 production. The high diffusivity of unbound PIP5K enables a broader range of inhibition, making it functionally more global than myosin-based negative feedback alone.

## MATERIALS AND METHODS

### Reaction-diffusion equations: STEN

The simulations are based on a previously described model in which three interacting species—RasGTP (Ras), PI(4,5)P_2_ (PIP2), and Protein Kinase B (PKB)—form an excitable network [16]. In this model, Ras and PIP2 are complementary: high Ras activity corresponds to low PIP2 levels and vice versa, a balance achieved through mutually inhibitory interactions that generate a positive feedback loop, characteristic of excitable networks [57]. PKB serves as the refractory species, activated by Ras but providing slow negative feedback. The concentrations of these molecules are governed by stochastic reaction-diffusion partial differential equations:

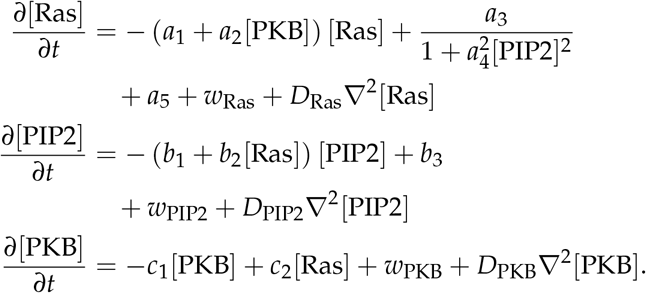

In each of these equations, the final term represents the diffusion of the species, where *D*∗ is the respective diffusion coefficient and ∇2 is the spatial Laplacian. The second-to-last terms represent molecular noise. Our model assumes a Langevin approximation in which the size of the noise is based on the reaction terms[68]. For example, in the case of PKB, the noise is given by

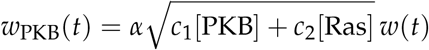

where *w*(*t*) is a zero-mean, unit-variance Gaussian, Brown-noise process. In the simulations, the size of this noise was adjusted with the empirical parameter *α*.

The system is largely robust to parameter changes, in that variations from the nominal values continue to give rise to excitable behavior [56]. To test this, we note that excitable behavior arises from the stochastic crossing of a threshold that is determined by the nullclines of the differential equations. To measure the level of robustness, we varied each of the parameters *a*_1_, …, *a*_5_, *b*_1_, …, *b*_3_, *c*_1_, and *c*_2_ from their nominal values and computed the size of the threshold. Some, like *a*_2_, *b*_2_, and *c*_1_, are quite robust, allowing large changes in either direction. Others allow large changes only in one direction.

Parameter values, given in Table 1, were selected to replicate experimental observations regarding excitable activities, such as the number of firings per unit time and wave propagation. Additionally, the choice was based on certain considerations. The diffusion was set such that the inhibitor diffused slightly faster than the activator. With these parameters, we observed wide waves that traveled large distances but did stop, and also occasionally we would get propagating waves, similar to those seen in Latrunculin-treated cells [69].

**Table 1.** STEN reaction-diffusion parameter values.

| Parameter | Value | Units |
| --- | --- | --- |
| $a_1$ | 0.0333 | $\text{s}^{-1}$ |
| $a_2$ | 0.7125 | $\mu\text{M}^{-1}\text{s}^{-1}$ |
| $a_3$ | 0.2850 | $\text{s}^{-1}$ |
| $a_4$ | 400 | $\mu\text{M}^{-1}$ |
| $a_5$ | $1.28 \times 10^{-4}$ | $\mu\text{M s}^{-1}$ |
| $b_1$ | 0.0036 | $\text{s}^{-1}$ |
| $b_2$ | 0.4920 | $\mu\text{M s}^{-1}$ |
| $b_3$ | 533.5 | $\mu\text{M}^{-1}\text{s}^{-1}$ |
| $c_1$ | 0.0186 | $\text{s}^{-1}$ |
| $c_2$ | 0.2160 | $\text{s}^{-1}$ |
| $D_F$ | 0.0075 | $\mu\text{m}^2\text{s}^{-1}$ |
| $D_B$ | 0.0075 | $\mu\text{m}^2\text{s}^{-1}$ |
| $D_R$ | 0.225 | $\mu\text{m}^2\text{s}^{-1}$ |
| $\alpha$ | 0.091 | |

### Reaction-diffusion equations: Polarization

In addition to the EN dynamics described above, we incorporate two additional terms related to cell polarization (see Fig. 1). The first of these feedback loops captures the influence of actin on polarity. Experimentally, increasing the assembly of branched actin has been shown to activate the Ras/PI3K signaling pathway, which is essential to promote the cell’s “front state,” leading to protrusion and movement [24]. Conversely, inhibiting branched actin results in an increase in cortical actin assembly and a strong suppression of Ras/PI3K activation, thereby promoting the ‘back-state’ of the cell. We represent the local abundance of branched actin by the variable *P*_*F*_. Furthermore, we assume that STEN regulates *P*_*F*_ through the concentration of PKB, as previous studies have shown that the synthetic recruitment of PKB to the membrane enhances actin polymerization [16]. Thus, we have:

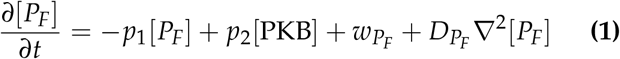

This component subsequently enhances Ras activity by modifying the production term of RasGTP:

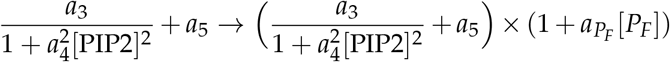

This establishes a positive feedback loop in which an increase in Ras leads to elevated PKB levels, which in turn enhances actin assembly, further promoting Ras production.

We also consider two inhibitory mechanisms. In the first, protrusions are assumed to increase tension, which consequently reduces Ras activity [55]. This model is similar to that of *P*_*F*_, but because tension acts globally, we represent it as follows:

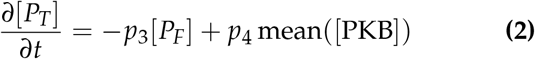

The mean value averages the concentration around the cell, representing a system in which the effect is global. Tension acts to inhibit Ras production, and we consider scenarios in which either the production of Ras is inhibited:

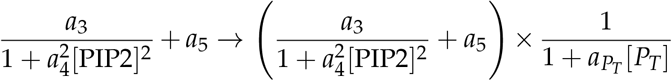

or its hydrolysis is enhanced:

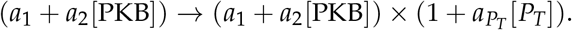

Because of the spatial nature of this negative feedback, we refer to this as the global-inhibition mechanism (see Fig. 1).

The second inhibitory pathway highlights the role of the acto-myosin network. Myosin II assembly leads to an immediate increase in Ras/PI3K activation, which enhances the cell’s sensitivity to chemotactic stimuli and promotes front-state behaviors, such as protrusion [24]. Conversely, myosin II disassembly reduces actin contraction at the cell’s rear, preventing the stabilization of the back-state. Together, this forms a feedback loop in which Ras/PI3K signaling promotes myosin II disassembly, and reduced myosin levels further enhance Ras/PI3K activation. We represent this interaction using a rear component, (*P*_*B*_), with its concentration given by:

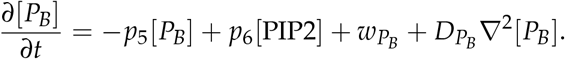

The input to this is PIP2, which represents a rear contribution [56]. These terms modify the components of the RasGTP equation related to hydrolysis:

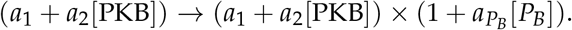

Note that, although this term is similar to that of *P*_*T*_, there are two main differences. First, this represents a local inhibition (see Fig. 1). More importantly, the feedback loop operates through the rear signal PIP2. Thus, while the net effect is to inhibit the front component (Ras), it achieves this by closing a positive feedback loop on the rear signal. Parameter values for these various feedback loops can be found in Table 2.

**Table 2.** Polarization loops parameter values.

| Parameter | Value | Units |
| --- | --- | --- |
| $p_1$ | 0.0125 | $\mu\text{M}^{-1}\text{s}^{-1}$ |
| $p_2$ | 0.0125 | $\mu\text{M}^{-1}\text{s}^{-1}$ |
| $a_{P_F}$ | 0.2–2.0 | $\mu\text{M}^{-1}$ |
| $D_{P_F}$ | 0.0075 | $\mu\text{m}^2\text{s}^{-1}$ |
| $p_3$ | 0.0125 | $\mu\text{M}^{-1}\text{s}^{-1}$ |
| $p_4$ | 0.0125 | $\mu\text{M}^{-1}\text{s}^{-1}$ |
| $a_{P_T}$ | 0.2–2.0 | $\mu\text{M}^{-1}$ |
| $p_5$ | 0.0125 | $\mu\text{M}^{-1}\text{s}^{-1}$ |
| $p_6$ | 0.0125 | $\mu\text{M}^{-1}\text{s}^{-1}$ |
| $a_{P_B}$ | 0.4–4.0 | $\mu\text{M}^{-1}$ |
| $D_{P_B}$ | 0.0013 | $\mu\text{m}^2\text{s}^{-1}$ |

### Simulating cell movement

To simulate cell movement, we adopted the procedure outlined previously [70]. In brief, PKB activity along the cell perimeter (in a one-dimensional simulation) was thresholded to generate a force normal to the cell surface. The vector sum of these forces around the perimeter was used to calculate a net force, which was scaled to fall within the range of experimentally observed protrusive stresses, 0.5-5 nN/*µ*m. This net force was then incorporated into a viscoelastic model of *Dictyostelium* mechanics [71].

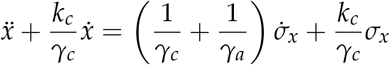

In this model, *σ*_*x*_ represents the *x*-component of the stress, with the viscoelastic parameters detailed in Table 3. A corresponding formula is used to denote displacement in the *y*-direction. The input to this linear viscoelastic model is derived from PKB activity, specifically *σ*_*x*_ ∝ [Ras]. The resultant displacements were normalized so that the maximum movement is equivalent to 10 *µ*m/min.

**Table 3.** Viscoelastic parameters [71].

| Parameter | Value | Units |
| --- | --- | --- |
| $\gamma_a$ | 6.09 | nN s/$\mu\text{m}^3$ |
| $\gamma_c$ | 0.064 | nN s/$\mu\text{m}^3$ |
| $k_c$ | 0.98 | nN/$\mu\text{m}^3$ |

#### Kymographs and heatmaps

For each model, the strengths of positive and negative feedback were varied from 0 to 20 in increments of 0.2. Each parameter set was simulated 20 times. PKB activity was visualized as kymographs, where the horizontal axis represents time and the vertical axis represents membrane activity. These kymographs were binarized, and activity patches were identified and analyzed using the regionprops function in MATLAB (MathWorks, Natick, MA). The following properties were measured: area, spatial extent, duration, and orientation. Patches were sorted in descending order based on area, and those smaller than 50 pixels were excluded from further analysis.

Heatmaps were generated to visualize the population-averaged trends in patch dynamics across parameter variations. The following metrics were analyzed:

1. **Mean patch width:** The kymographs were binarized to identify patches and MATLAB’s BoundingBox command was used to determine the spatial and temporal boundaries of each patch. The spatial width was averaged across all patches within each simulation, and the mean values were plotted as a heatmap.
2. **Duration of the longest patch:** The temporal extent of each patch was extracted using BoundingBox. The duration of the largest patch (by area) within each simulation was recorded, and the average across simulations was visualized.
3. **Patch frequency:** To assess persistent activity patterns, transient patches were filtered out. At each time point, the number of patches present on the membrane was counted. The most frequently observed patch count over the simulation defined the kymograph’s dominant state, which was then mapped across parameter space.
4. **Polarization duration:** A cell was considered polarized when exactly one membrane patch was present. The percentage of simulation time spent in this state was recorded and visualized as a heatmap.

### Determination of polarized cells

A cell was classified as polarized if it met the following criteria:

- The most frequent number of membrane patches throughout the simulation was one.
- The cell spent more than 10% of the total simulation time in the polarized state.

To exclude hyperactive cells with a single, large patch covering most of the membrane, cells with a patch orientation angle exceeding 60° were excluded. The total number of simulations meeting these criteria was reported as the number of polarized cells.

### *Dictyostelium* experiments

*Dictyostelium* cells were used to observe signaling activity (Fig. 2) and quantify movement (Fig. 5).

#### Cells and plasmids

*Dictyostelium discoideum* cells were cultured in HL5 media for a maximum of 2 months after thawing from frozen stock. AX3 cells were obtained from the R. Kay laboratory (MRC Laboratory of Molecular Biology, UK).

Plasmids were introduced into *Dictyostelium* cells using electroporation. To improve efficiency, heat-killed *Klebsiella aerogenes* was added after transformation. pDM358 RBD-EGFP, based on the 51–220 amino acids of Raf, was previously created in the Devreotes lab.

#### Microscopy

Cells were seeded in 8-well Lab-Tek chambers (Thermo-Fisher, 155409) and left to settle for ten minutes before the media was gently aspirated and replaced with Development Buffer (DB) (5 mM Na_2_HPO_4_, 5 mM KH_2_PO_4_, 1 mM CaCl_2_, 2 mM MgCl_2_). Cells were allowed to sit for an hour starving in DB prior to imaging to decrease photosensitivity. For experiments with polarized cells, growth-phase cells were washed and suspended in DB at a density of 2 × 107 cells/ml. Cells were then developed by being shaken for 1 hour and subsequently pulsed with 50–100 nM cAMP every 6 minutes and shaken for 4 hours.

Laser scanning confocal imaging was carried out on two microscopes: a Zeiss AxioObserver inverted microscope with an LSM800 confocal module and a Zeiss AxioObserver with 880-Quasar confocal module and Airyscan FAST module. On the LSM800 microscope, GFP and YFP proteins were excited with a solid-state 488 nm laser and on the 880 with an argon laser. Emission wavelengths collected were chosen to avoid overlap between GFP and mCherry emission profiles. All imaging was done with 63 × /1.4 PlanApo oil DIC objectives and appropriate raster zoom. Brightfield images were acquired using a transmitted-photomultiplier tube (T-PMT) detector.

Spinning disk confocal imaging was performed on a Nikon TiE2 CSU W1 SoRA microscope, a solid-state 488 nm laser, a 60 × /1.49 Apo TIRF objective, and a Hamamatsu FusionBT camera. Camera binning was adjusted to optimize for signal-to-noise ratio, resolution, and speed of acquisition. All images for a given experiment were acquired at the same binning.

#### Preparation of reagents and inhibitors

For experiments with latrunculin (Fig. 2), 2.5 *µ*l aliquots of 1 mM latrunculin in DMSO (Millipore Sigma, 428026) were diluted 1:20 in DB to make a 10 × (50 *µ*M) stock. 50 *µ*l of this stock was then added to a well containing 450 *µ*l of media prior to imaging. Cells were incubated in latrunculin for at least 10 minutes prior to the start of the experiment.

#### KikGR optogenetic florescence to study dynamic partitioning of PIP5K

PIP5K (phosphatidylinositol 4-phosphate 5-kinase) photoconversion can be achieved using KiKGR, a green-to-red photoconvertible fluorescent protein. KiKGR is initially fluorescent green when excited with 488 nm light. Upon exposure to blue and near-ultraviolet light (405nm), it undergoes an irreversible photoconversion, shifting its emission from green (516 nm) to red (593 nm). This property is used to mark specific regions or populations of proteins in living cells and then track their movement over time.

PIP5K is genetically fused to KiKGR, allowing the fluorescent tag to report on the localization and dynamics of the kinase. Before photoconversion, the fusion protein fluoresces green, highlighting the regions where PIP5K is localized (often at the plasma membrane or specific intracellular compartments). A targeted area of the cell is exposed to 405 nm light using a laser-scanning confocal microscope. This exposure converts KiKGR from green to red in the illuminated region, without affecting unexposed areas. After photoconversion: The red fluorescence represents the PIP5K molecules present at the moment of photoconversion. The green fluorescence represents newly synthesized or newly trafficked PIP5K molecules.

By imaging both channels over time, you can track: PIP5K turnover rates from how fast the green signal fades as new red protein appears; protein trafficking from movement of red-labeled PIP5K from one cellular region to another; and diffusion dynamics, if the photoconverted region spreads over time.

#### Cell tracking and classification

Cell tracking in Fig. 5 was performed using the TrackMate plugin in ImageJ. The plugin employs an intensity thresholding approach to distinguish the foreground from the background and connects regions with similar intensities within enclosed boundaries. For each identified region, several properties were measured, including mean intensity, orientation, circularity, and the position of the centroid. To track the trajectories of individual cells over time, the Linear Assignment Problem (LAP) tracker was utilized, allowing for accurate linking of detected spots across sequential frames. The mean circularity values obtained from the segmentation process were further used to classify cells into unpolarized, polarized, and intermediate polarized states. This classification was based on a negative correlation observed between cell speed and circularity, determined by linear regression analysis.

#### Kymograph generation for membrane activity of live cells

Kymographs representing membrane activity along the cell perimeter over time were generated from binarized image stacks of individual cells. Inner and outer radii were defined relative to the cell centroid using the bounding box properties from the regionprops function in MATLAB, ensuring coverage of the entire cell boundary. Membrane intensity (IM) at equally spaced points along the perimeter was quantified by performing 1-pixel-wide linescans perpendicular to the boundary. The length of each linescan was adjusted based on the defined inner and outer radii to ensure complete coverage of the boundary across all angles. For each linescan, IM was calculated as the average of the three brightest pixels. These intensity values were plotted as a function of time and perimeter angle to construct the kymograph.

### HL-60 experiments

Neutrophil-like HL60 cells were used to observe the effect of various pharmacological interventions on cell activity (Fig. 7).

#### Reagents and inhibitors

200 *µ*g/ml fibronectin stock (Sigma-Aldrich; Cat #F4759-2MG) was prepared in sterile water, and subsequently diluted in 1x PBS. 10 *µ*M Latrunculin B (Sigma-Aldrich; Cat #428020) or 50 mM CK666 (EMD Millipore; Cat #182515) stock was prepared in dimethyl sulfoxide (DMSO, Sigma-Aldrich; Cat #D2650). Phorbol 12-myristate 13-acetate (PMA, Sigma-Aldrich; Cat #P8139) was dissolved in DMSO to make 1 mM stock. Puromycin (Sigma-Aldrich; Cat #P8833) or blasticidin S (Sigma-Aldrich; Cat #15205) was prepared in sterile water to make stock solution of 2.5 mg/ml or 10 mg/ml, respectively. Aliquots of all stock solutions were stored at ™ 20 °C. According to experimental requirements, further dilutions were prepared in 1 × PBS or culture medium before adding to cells.

#### Cell culture

Female human HL-60 cells were grown in RPMI 1640 medium (Gibco; Cat #22400-089) supplemented with 15% heat-inactivated fetal bovine serum (Thermo Fisher; Cat #16140071) as described previously [72, 73]. To obtain migration-competent neutrophils or macrophages, WT or stable lines were differentiated in the presence of 1.3% DMSO over 6–8 days or 32 nM PMA for 2–3 days, respectively [56, 72, 73]. All cells were maintained in humidified conditions at 5% CO_2_ and 37 °C. Stable cells were maintained in the presence of selection antibiotics, which were removed during differentiation and experimentation.

#### Plasmid construction and transfection

All DNA oligonucleotides were procured from Sigma-Aldrich. CRY2PHR–mCherry–KRas4B G12VΔCAAX/pPB (Addgene #201753), CIBN–CAAX/pLJM1 (Addgene #201749) or LifeAct–miRFP703/pLJM1 (Addgene #201750) construct was generated in a previous study [73]. DNA sequence (996 bases) encoding the 5-phosphatase, INP54P, was PCR-amplified and subcloned into the BspEI and SalI sites of the PiggyBac transposon plasmid to create CRY2PHR-mCherry-INP54P/pPB construct. All constructs were verified by diagnostic restriction digestion and sequenced at the institutional sequencing facility.

HL-60 cells stably co-expressing CIBN-CAAX, LifeAct-miRFP703, and opto-KRas4B G12VΔCAAX were constructed earlier [73]. Here, we introduced 5 *µ*g opto-INP54P construct with an equal amount of PiggyBac transposase plasmid in the CIBN-CAAX and LifeAct-miRFP703 co-expressing HL-60 stable cell line using Neon transfection system 100 *µ*l kit (Thermo Fisher; Cat #MPK10025). Cells were selected in the presence of puromycin and blasticidine S.

#### Confocal microscopy and optogenetics

Differentiated neutrophils or macrophages were adhered on an eight-well coverslip chamber in the presence or absence of fibronectin, respectively. Next, fresh RPMI 1640 medium was added to attached cells and used for imaging. All imaging was performed at a middle plane of the cells with 0.2–10% laser intensity using a Zeiss LSM780-FCS single-point, laser scanning confocal microscope supported with ZEN Black software. Images were acquired with a 40 × /1.30 PlanNeofluar oil DIC objective, in addition to digital zoom. For inhibitor experiments, cells were treated with 50 µM CK666 or 5 µM latrunculin B for at least 10 minutes before imaging [56, 72, 73].

Optogenetic experiments were done in the absence of any chemoattractant, as described previously [56, 72, 73].

Briefly, a solid-state laser (561 nm excitation and 579– 632 nm emission) was used for visualizing opto-KRas4B G12VΔCAAX or opto-INP54P whereas a diode laser (633 nm excitation and 659–709 nm emission) was utilized to capture LifeAct-miRFP703 expression. Images were acquired for approximately 5 minutes, after which a 450/488 nm excitation laser was turned on globally to activate INP54P or KRas4B G12VDCAAX recruitment. Image acquisition and photoactivation were done at 7-second intervals.

## ACKNOWLEDGMENTS

We thank members of the Iglesias and Devreotes labs for useful discussions. Special thanks to Bedri Abubaker-Sharif for valuable insights on the computational models and Yiyan Lin for sharing her experimental results. Additional acknowledgment to Doug Robinson (JHU School of Medicine) for his feedback and insight.

