## Supplementary figures and images for "Spatial distribution of cytoskeleton-mediated feedback controls cell polarization: a computational study"

### S1 Figure

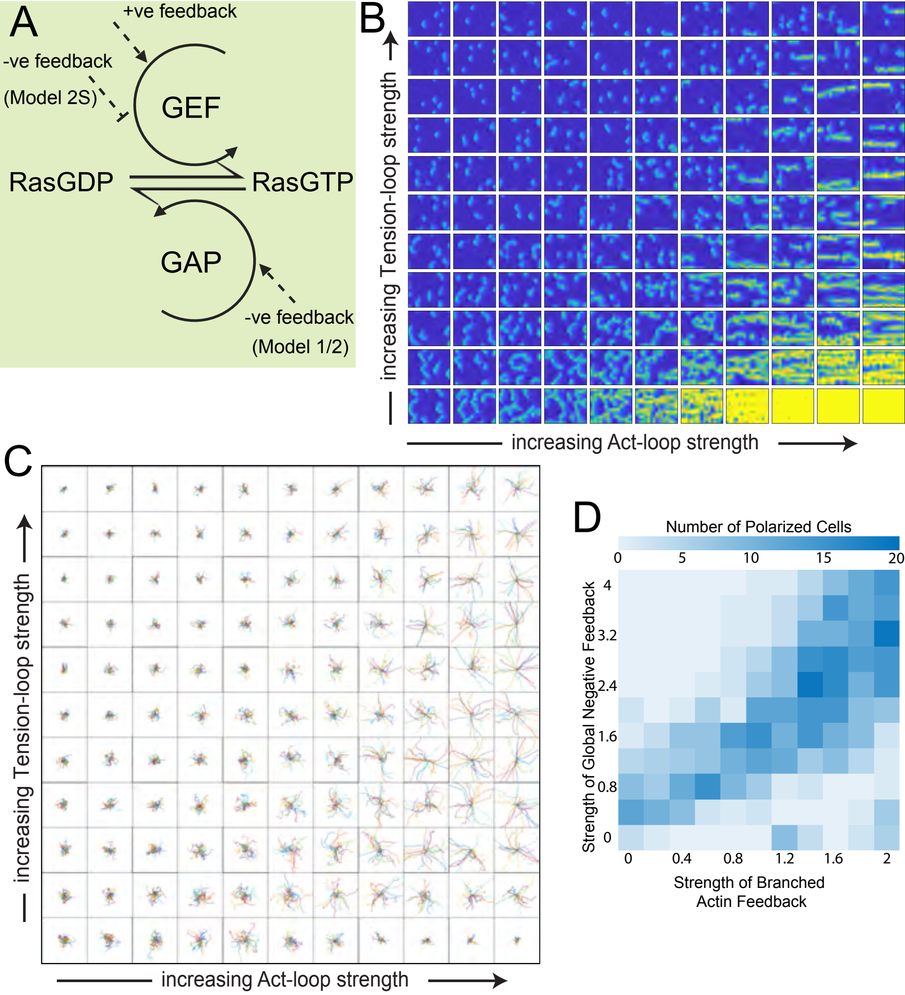

### S2 Figure

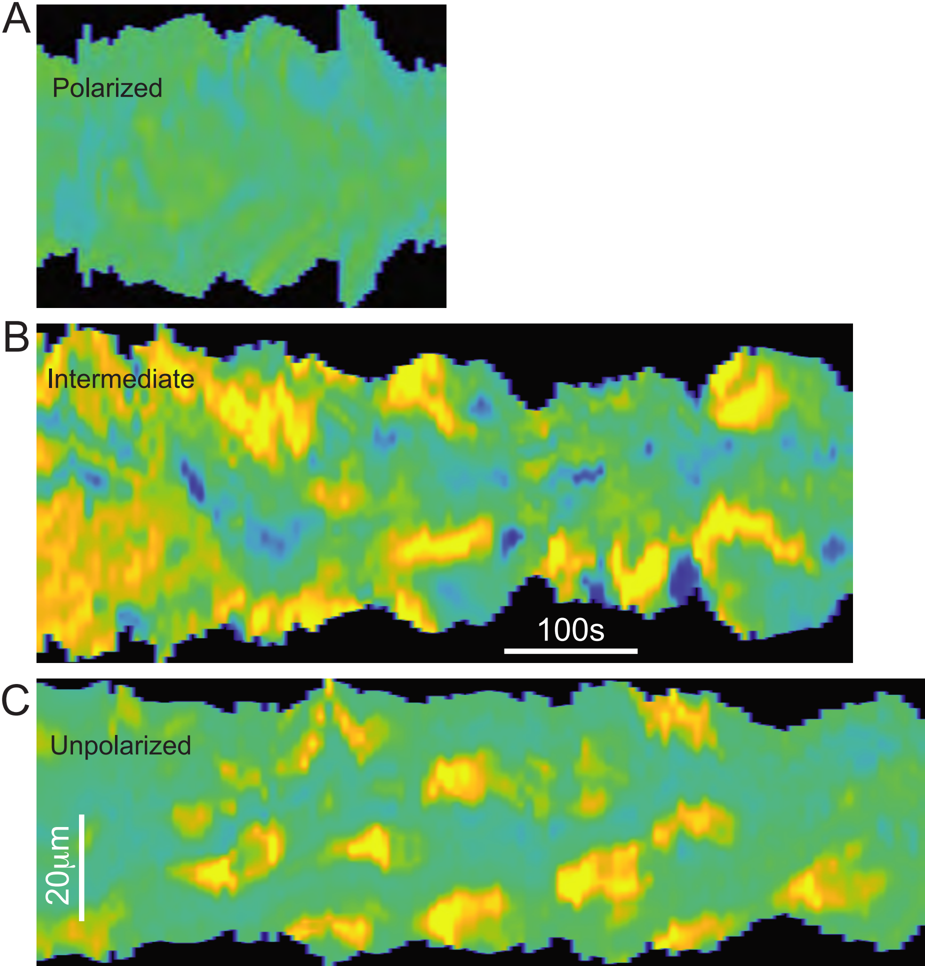

### S3 Figure

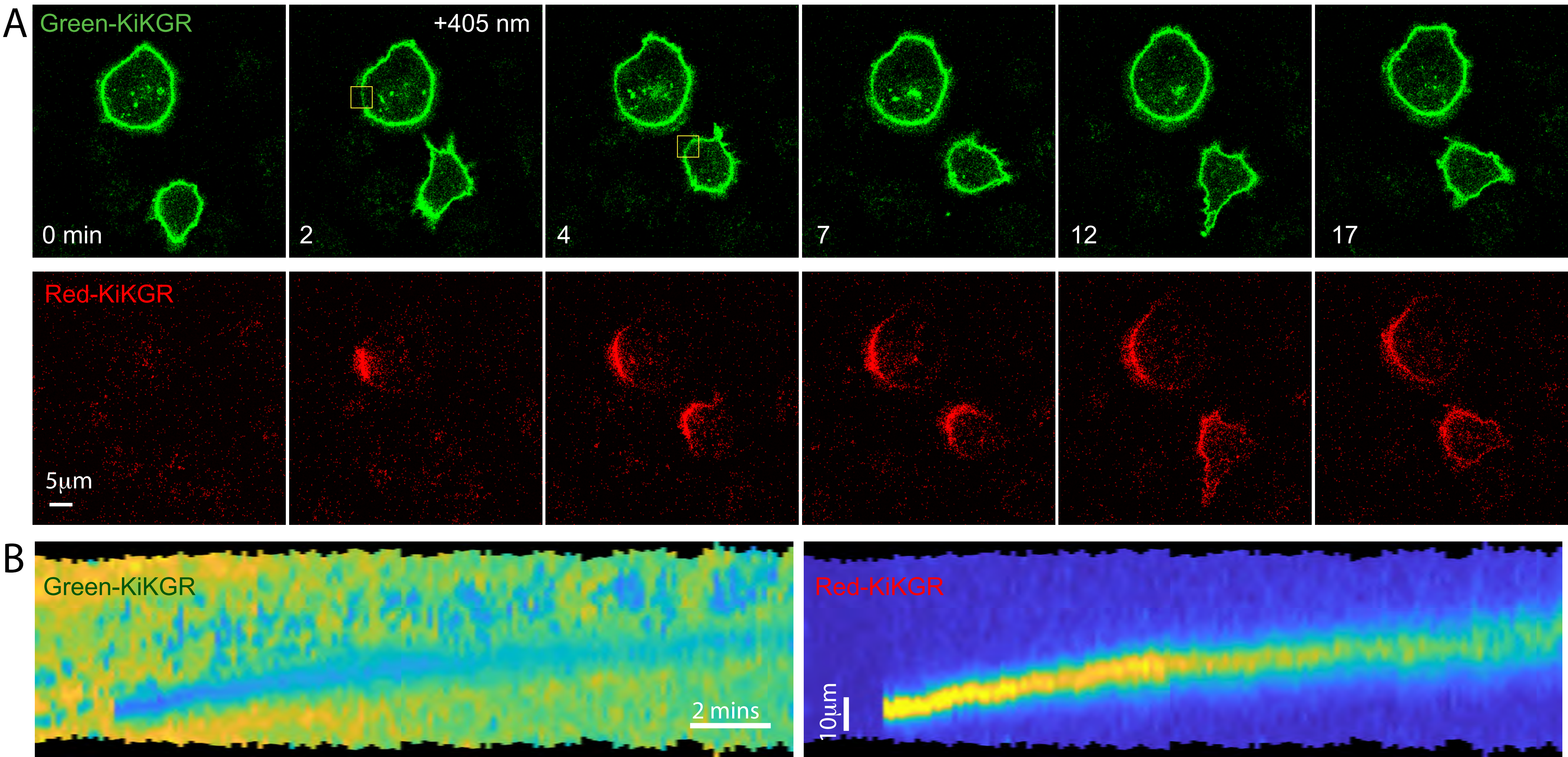
